# Cannibalism as a feeding strategy for mantis shrimp Oratosquilla oratoria (De Haan, 1844) in the Tianjin coastal zone of Bohai Bay

**DOI:** 10.1101/740100

**Authors:** Qi-Kang Bo, Yun-Zhao Lu, Hui-Jing Mi, Yan-Guang Yu, De-Xian Gu, Hong-Zheng You, Shuang Hao

**Affiliations:** Breeding Station of Bohai Sea Aquatic Resources, Tianjin Fisheries Research Institute; Marine Resources and Ecology Research Department, Tianjin Fisheries Research Institute; Aquatic Disease Research Department, Tianjin Fisheries Research Institute

**Author notes:** Corresponding Author (Y-G Y).

## Abstract

A representative semi-enclosed bay of China, Bohai Bay has experienced severe interference in recent decades and is under threat from rapid human development. Although the mantis shrimp *Oratosquilla oratoria* plays an important role in the ecosystem and fishery, its feeding ecology and the impact of habitat changes on its feeding habits are poorly known. In this study, we sought to identify the prey consumed by *O. oratoria* through the separation of stomach contents and to describe its trophic ecology during maturation, from March to July, in the Tianjin coastal zone of Bohai Bay. A total of 594 specimens were collected and 347 (58.59%) stomachs were found to have food remains. More than half of the *O. oratoria* individuals had poor feeding activity, and the degree of feeding activity of females was higher than that of males, but there was no significant difference in the visual fullness index and the fullness weight index (FWI) between sexes for each month. And the feeding activities of O. oratoria were consistent over the study months. A total of 207 prey items yielded 231 readable sequences and 24 different taxa were identified. Prey detected in *O. oratoria* consisted mainly of crustaceans, which accounted for 71.86 % of the clones detected; 16.02% corresponded to fishes, 8.23% corresponded to mollusks and the remaining 3.90% corresponded to other marine organisms. Cannibalism (occured frequently, 69.08%) in this study was noticeably higher than that seen in previous studies and confirmed that cannibalism may be a significant feeding strategy in the mantis shrimp *O. oratoria* in the Tianjin coastal zone of Bohai Bay. The ecological environment in Bohai Bay has been affected by anthropogenic activities and the macrofaunal biodiversity and abundance have noticeably declined, which might make the food scarce for the mantis shrimp *O. oratoria*. Then, the starvation obviously increased cannibalistic tendencies.

## Introduction

The mantis shrimp, *Oratosquilla oratoria* (De Haan, 1844) (order Stomatopoda) is well known as a ferocious predator with its large and powerful raptorial appendages. It is a benthic, neritic and burrowing shrimp that is found on muddy bottoms in the coastal waters of Siberia, Korea, China, Japan, Vietnam, and Australia [1, 2]. It has become a commercially important species in these regions.

In Bohai Bay, the mantis shrimp is heavily caught by bottom-trawl and trammel nets, whose annual catches account for more than one-third of the crustacean catches in the past ten years [3]. A substantial decrease in the stock size of large female shrimps has been apparent since the fishing industry catch, in which larger individuals are overexploited, and the season of spawning fastigium is delayed to mid-to-latter of May, while some female shrimps also spawn into September [1, 4–6]. The abundance of mantis shrimp pseudozoea is low before July [4, 6].

Few studies have focused on the feeding ecology of *O. oratoria*. Its trophic ecology in the coastal waters of the Chinese open sea, Huanghai Sea and Donghai Sea was described with crustaceans being the main prey group, followed by fishes and polychaetes [7, 8]. Hamano et al (1986) [9] found that mantis shrimp feed largely on Macrura and Pelecypoda in Hakata Bay of Japan. However, the diet of marine organisms is affected by prey availability and the composition of marine life in the surrounding environment [9, 10]. In other distribution areas, such as Bohai Bay, their ecology is poorly known. Bohai Bay is a typical semi-enclosed bay situated in the western part of the largest inner sea of China, Bohai Sea which is located in northern China. As one of the most important marine fisheries and natural resource reserves in China, the ecological environment of Bohai Sea has always been of wide concern [11, 12]. In Bohai Bay, mantis shrimp, a very intensive predator, is a principal component of the benthic megalofaunal community that consumes and transfers energy and biomass from the base of the web to higher levels [10, 13]. Moreover, an understanding of the trophic interactions between prey and predators is the foundation for effective management and protection of natural resources [14].

Intensive anthropogenic activities have been exerting tremendous stress on marine organisms, resulting in significant changes and deterioration in the structure of biological communities [11, 15]. The study area, the Tianjin coastal zone of Bohai Bay, has experienced rapid economic and technological development and anthropogenic activities such as terrigenous pollution, aquaculture, transportation and offshore oil exploration have caused serious impacts on the coastal environment [12, 16]. However, how the impact of habitat changes on the feeding habits of a commercially important species, mantis shrimp, is unknown.

Artificial breeding of mantis shrimp has been explored, and the promotion of gonadal development has become a knotty technical problem in artificial cultures [17]. In the absence of available formulated bait, the discovery of the feeding ecology during gonad maturation will provide a reference for aquaculture and artificial breeding of this commercially important species.

Because of the importance of this species to the ecosystem and for fisheries, we sought to identify the prey consumed by *O. oratoria* through the separation of stomach contents, and to describe its trophic ecology during maturation in the Bohai Bay using the DNA barcoding method.

## Material and methods

### Study area and samples acquisition

Tianjin city is situated in the western part of Bohai Bay in China, and the study area was the Tianjin coastal zone (Fig 1). All the analyses have been carried out using frozen dead specimens of mantis shrimp collected from local fishermen from March to July 2018. Samples of *O. oratoria* were obtained from Bohai Bay by bottom trawls which were conducted by pleasure-boats that were converted from fishing boats, and on boats part of the catch was cooked on the spot for amusement and food; the rest of the catch was frozen and then distributed to tourists at the end of the tour. No use of live animals has been required for this study and no specific permissions were needed for the sampling activities in all of the investigated areas because our species of interest is commercially harvested (not endangered nor protected) and was caught in areas where fishing is allowed.

**Fig 1.**
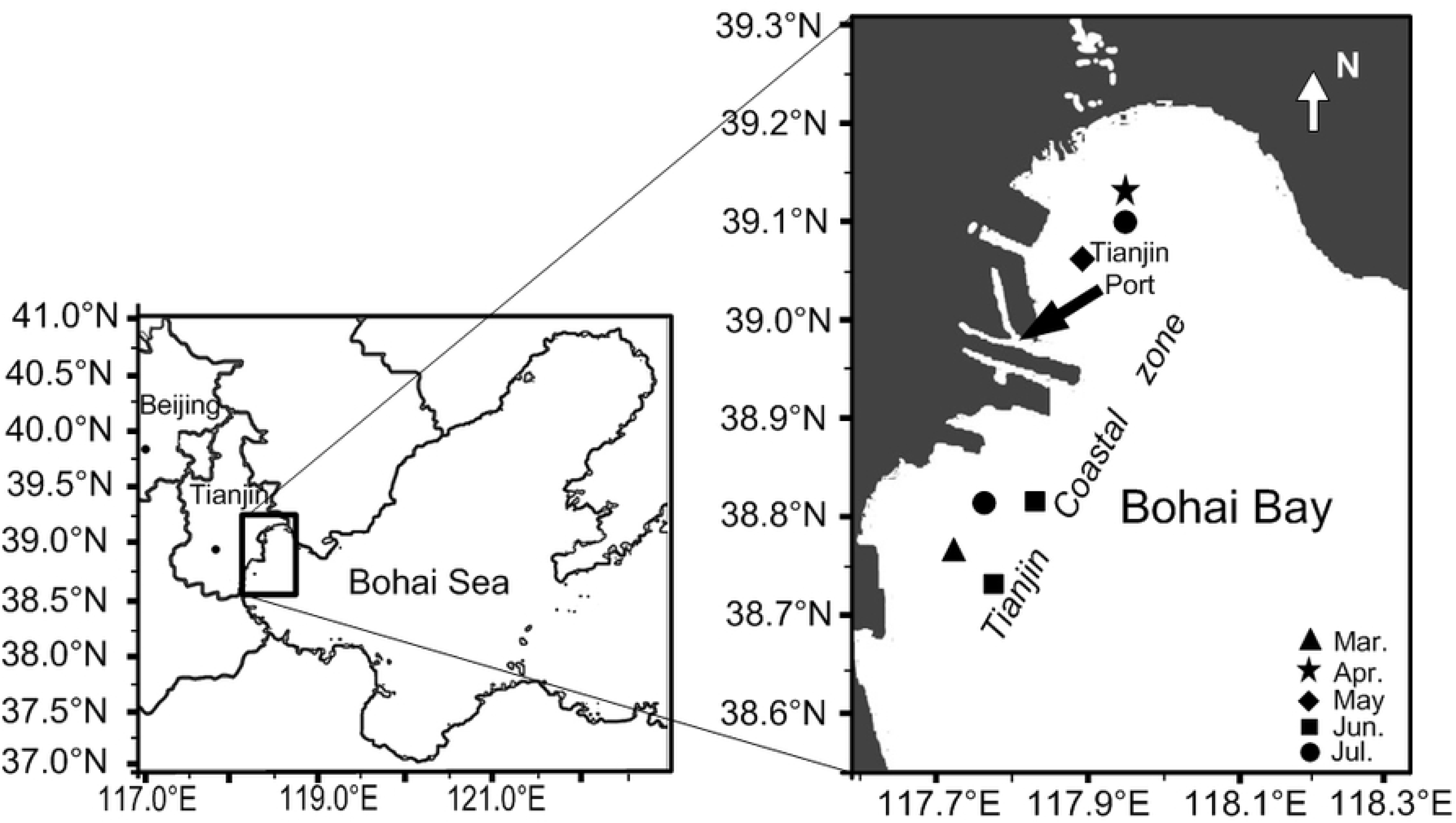
The location of the sampling sites in the Tianjin coastal zone of Bohai Bay; the marks show the sampling sites. Marks of each shape represent sampling sites in different months.

All specimens were brought to the laboratory where their stomach contents were removed. Each stomach was opened and the contents were flushed into cryogenic vials. The stomach contents, potential prey residue, were weighed and then preserved in 70% ethanol at −20℃ for later DNA analysis. To avoid potential contaminants (e.g., blood and tissue attached to the stomach from the predator), the exterior surface of each stomach was washed with sterile, distilled water before removing the stomach contents [18]. The total length (TL, from the base of the eyestalk to the anterior edge of the median notch of the telson) of the shrimps was measured to the nearest 1 mm, stomach content weight (SCW) and the total body weight (TW) were obtained to the nearest 0.001 g. This basic information and collection dates were showed in Table 1.

**Table 1.** The collection date, total length (TL), total weight (W), number (N) of *O. oratoria* specimens and number of stomach content specimens (Nsc) obtained from the Tianjin coastal zone of Bohai Bay.

| Date | TL (cm) | W(g) | N | Nsc |
| --- | --- | --- | --- | --- |
| 17-Mar | 12.30±1.53<sup>a</sup> | 25.391±9.95<sup>a</sup> | 34 | 11 |
| 20-Apr | 12.15±2.16<sup>a</sup> | 21.670±7.35<sup>ab</sup> | 155 | 86 |
| 14-May | 11.66±1.28<sup>ab</sup> | 19.923±6.34<sup>b</sup> | 110 | 67 |
| 10, 25-Jun | 10.48±1.12<sup>b</sup> | 11.923±3.50<sup>c</sup> | 150 | 84 |
| 16, 29-Jul | 12.29±1.63<sup>a</sup> | 21.609±5.19<sup>ab</sup> | 145 | 99 |

Data with different letters significantly differ ( $p < 0.05$ ) among months.

Feeding intensity during the study months was determined based on the degree of fullness of the stomach. A visual stomach fullness index was assigned to 5 levels: 0= empty, 1= scarce remains, 2= half full, 3= almost full, and 4= completely full [19]. The mantis shrimp with stomachs at level 2 to 4 were considered to have been actively feeding, while stomach at 0 and 1 levels were considered to indicate poor feeding activity, and the percentages of shrimps with actively feeding (AF) and poor feeding activity (PFA) in both sexes for each month were calculated. The fullness weight indices (FWI= (SCW×1,000)/TW) were calculated for all mantis shrimp with food remains [20].

### DNA extraction and sequence acquisition

Stomach contents were evenly ground in a homogenizer. Given few items and a large amount of silt in the stomach contents sample, genomic DNA was isolated using Soil Genome DNA Extraction Kit DP336 (TIANGEN BIOTECH, CO., LTD). A fragment of the *mitochondrial cytochrome oxidase I* (CO I) gene was amplified using the universal primers LCO1490-HC02198 [21] and if the concentration of amplification productions did not meet the sequencing requirements, then a semi-nested PCR using universal primers mlCOIintF-jgHCO2198 [22] was performed to increase the copies of prey DNA.

Each polymerase chain reaction (PCR) was carried out in 50-µL volumes containing 2U Taq DNA polymerase (Takara Co.), approximately 20 ng template DNA, 0.2 mM dNTPs, 0.25 µM of each primer, 2.5 mM MgCl_2_ and 1×PCR buffer. The PCR amplification was performed on a GeneAmp® 9700 PCR System (Applied Biosystems). Cycling conditions consisted of an initial denaturation at 94℃ for 3 min, followed by 35 cycles of: denaturation at 94℃ for 1 min, annealing at 50℃ (54℃ for primers mlCOIintF-jgHCO2198) for 30s, and extension at 72℃ for 45s, and a final step of 5 min at 72℃. The semi-nested PCR was carried out using 1µL of the first PCR as a template.

Amplification products were confirmed by 1.5% TBE agarose gel electrophoresis stained with ethidium bromide. The cleaned product was prepared for sequencing using the BigDye Terminator Cycle Sequencing Kit (ver.3.1, Applied Biosystems) and sequenced bidirectionally using an ABI PRISM 3730 (Applied Biosystems) automatic sequencer. Obtained sequence producing mixed peak indicated that more than one prey species were present. Those PCR products were cloned using the TOPO TA Cloning Kit (Invitrogen). Eight colonies per sample were selected for colony PCR amplification and sequencing using the primers M13 (forward): GTAAAACGACGGCCAG, andM13 (reverse): CAGGAAACAGCTATGAC.

### DNA analyses

All the obtained sequences were assembled and edited separately using DNASTAR software (DNASTAR, Inc.) and were then submitted and identified using the Identification System (IDS) in the Barcode of Life Database (BOLD, www.boldsystems.org) and the Basic Local Alignment Search Tool (BLAST) query algorithm in GenBank to establish whenever possible the identification of the ingested material. The criteria to assign identification at the species level required that the sequence similarity display >98% in the BOLD database or in BLAST [23]. When a similar sequence match was not found in the DNA barcode reference library, we applied the method for visualization of a neighbor-joining tree and based our taxonomical assignments following the strict criteria proposed, and consist in nesting the ‘‘unknown’’ within a clade comprising of members of a single taxon [24]. The neighbor-joining tree was constructed using MEGA 6 software based on the Kimura 2-parameter model [25] and bootstrap probabilities with 1,000 replications were calculated to assess reliability on each node of the tree.

### Statistical analysis

To corroborate that the number of analyzed stomachs was adequate for diet description, a cumulative prey curve was generated using Estimate S Version 8.2 based on the prey identified. The number of samples was assumed to be sufficient to describe the diet when the curve approached the asymptote [26]. Differences in mean TL and TW between months and in the FWI of males and females were compared using Student’s t-test. To look for significant differences in FWI in different months for males and females, one-way analysis of variance (ANOVA) was conducted. Homogeneity of variance was examined for all data by using Bartlett-Box F and Cochran’s C tests. In this study, the distribution of FWI values was not uniform for each month and the data did not meet the assumptions of normality and homogeneity of variances (P<0.05). Therefore, the FWI was transformed into a log (x) scale to normalize and homogenize the variances [27]. The distribution differences of AF and PAF between sexes were verified by the Chi-squared fit test (χ^2^). Statistical analyses were performed using IBM SPSS statistics version 19 (IBM, Chicago, IL, USA). The frequency of occurrence of each prey was calculated as the percentage of stomachs in which the prey occured in any given sample. The number (%N) was calculated as the number of a certain prey type relative to the total numbers of prey.

## Results

### Feeding intensity

A total of 594 specimens were collected and 347 stomachs (58.59% of total stomachs) were found to have food remains. More than half of the stomachs were found to have low index values (the mean proportion of PAF in females was 68.86%, and in males was 75.71%). Although the analysis of the Chi-squared fit test show that there was no significant difference in the visual fullness index between sexes (P>0.05), the AF proportions of females were larger than those of males except in June (Fig 2).

**Fig 2.**
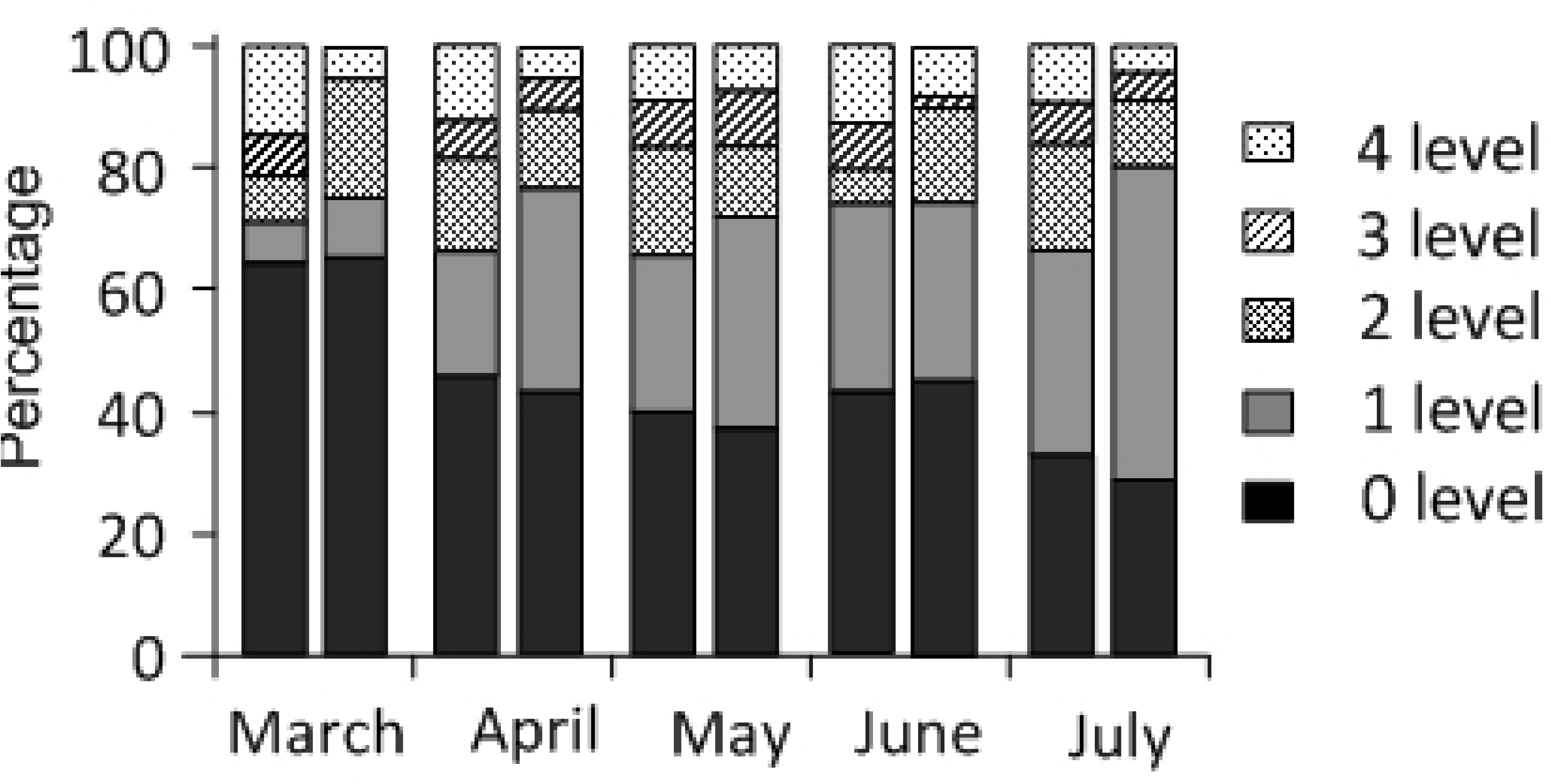
Monthly variation in fullness degree of stomachs of *O. oratoria* for each month. The first column shows data for females and the second column shows data for males.

From March to July, most FWI values for the females ranged from 1.54 to 7.52 (Fig 3a) which were larger than those for males, from 1.49 to 4.83 (Fig 3b), but no difference was found in the FWI between sexes for each month or between months for each sex, after comparison with the transformed data (P>0.05).

**Fig 3.**
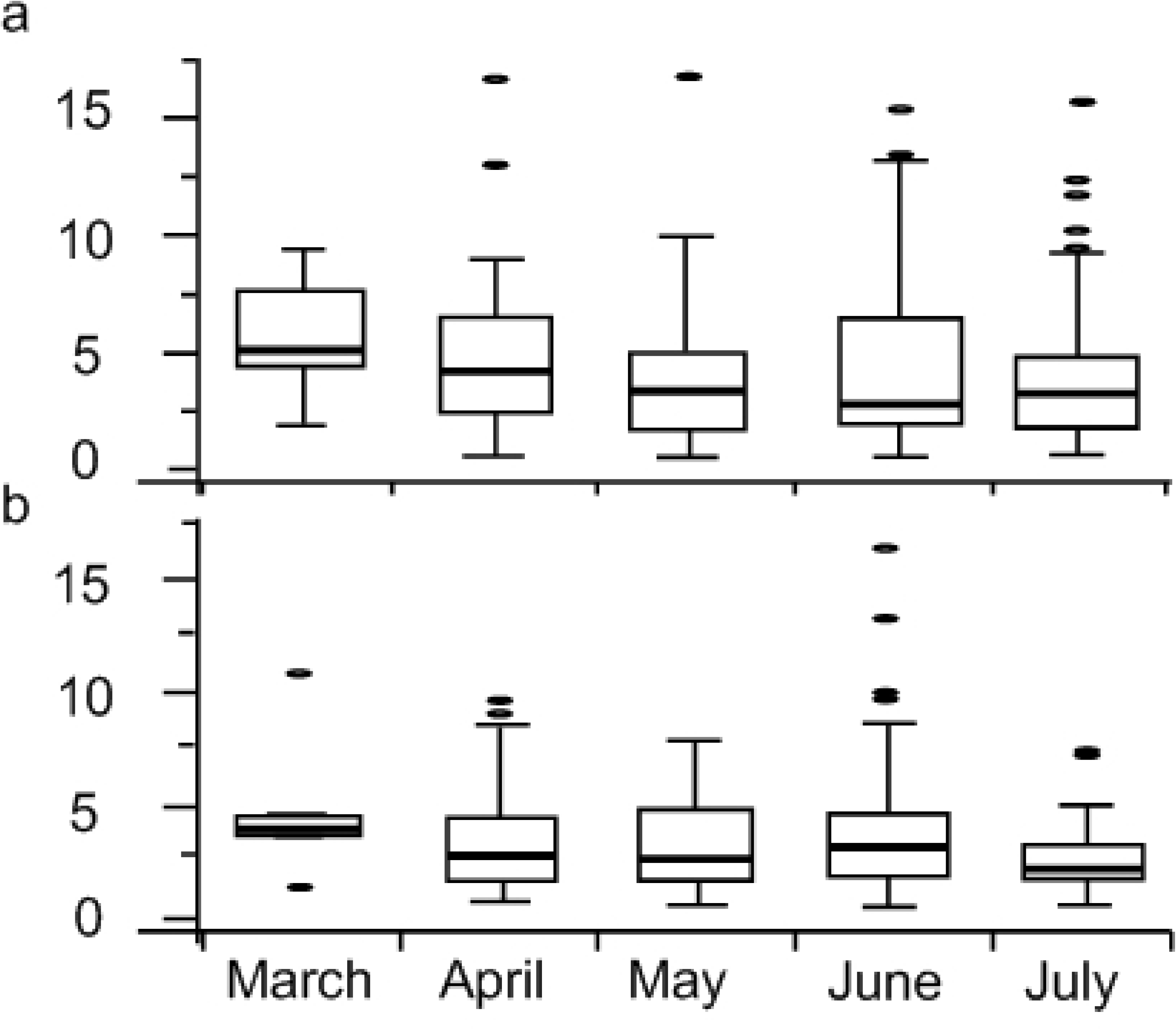
Box-plots of the fullness weight index (FWI) of the females (a) and males (b) in different months. Boxes show lower and upper quartiles with medians (lines) inside the boxes.

### Molecular prey identification

For 347 prey items obtained, about 95% contained a few items and a large amount of silt. Except for a large number of exopods or endopods remains of pleopods of mantis shrimps and a small amount of skeletal fragments from fish, there was no hard skeletal material in the stomach contents. All prey tissue fragments were barcoded, and 207 pray items yielded 231 readable sequences. The sequences were compared to the sequences of the reference library, and 90.91% matched with greater than 98% similarity to the reference sequences, allowing identification to the species level. The remaining 9.09% (21 clones) could be identified only to the genus, family or order level (Table 2). Of the 21 clones, only one clone showed more than a 95% similarity to reference sequences, while the remainder showed similarities between 82 and 91%. A total of 24 different taxa were identified. The representative sequences were submitted to GenBank (submission ID is: 2250197) and also submitted as Supporting information (S1 File).

**Table 2.** Prey detected in *O. oratoria* stomachs from Bohai Bay by cloning the CO I fragment gene, including GenBank Accession numbers or Sequence IDs of closest matches, percentages of similarity obtained from BLAST and BOLD, prey taxa and their monthly frequencies.

| Order | Family | species | Similarity % | Ac. Number or<br>Seq. ID | Frequence |  |  |  |  |
| --- | --- | --- | --- | --- | --- | --- | --- | --- | --- |
|  |  |  |  |  | Mar. | Apr. | May | Jun. | Jul. |
| Gobiiformes | Oxudercidae | *Chaeturichthys stigmatias* | 100 | KV199164 | 1 | 2 | 11 | 10 | 6 |
| Gobiiformes | Oxudercidae | *Odontamblyopus rubicundus* | >98 | AF391371 |  |  |  | 2 | 1 |
| Gobiiformes | Oxudercidae | *Amblychaeturichthys hexanema* | 98.6 | GU479054 |  | 1 |  |  |  |
| Clupeiformes | Clupeidae | *Konosirus punctatus* | 99.28 | JQ753955 |  |  |  |  | 1 |
| Clupeiformes | Engraulidae | *Thyssa kammalensis* | 98.12 | KU360510 |  |  |  |  | 1 |
| Scorpaeniformes | Platycephalidae | *Platycephalus indicus* | 98.1 | JN885883 |  | 1 |  |  |  |
| Stomatopoda | Squillidae | *Oratosquilla oratoria* | >99 | KP976321 | 7 | 47 | 29 | 29 | 31 |
| Mysida | Mysidae | No match found |  |  |  | 3 | 6 | 2 | 2 |
| Decapoda | Hippolytidae | No match found |  |  |  | 1 |  |  |  |
| Decapoda | Alpheidae | *Alpheus distinguendus* | 98.39 | GQ892049 |  |  |  |  | 1 |
| Decapoda | Dorippidae | *Heikea arachnoides* | 99.27 | EU636976 |  |  |  |  | 2 |
| Decapoda | Porcellanidae | No match found |  |  |  |  |  |  | 1 |
| Decapoda | Upogebiidae | Austinogebia | 95.61 | LC006054 |  |  | 1 |  |  |
| Decapoda | Varunidae | No match found |  |  |  |  |  |  | 3 |
| Amphipoda | Amphipoda | No match found |  |  |  |  | 1 |  |  |
| Myopsida | Loliginidae | *Loliolus beka* | >99 | HQ529504 |  |  | 3 | 1 | 8 |
| Octopoda | Octopodidae | *Octopus minor* | 99.55 | MF029677 | 2 |  |  |  |  |
| Littorinimorpha | Naticidae | *Neverita didyma* | 99.51 | JF693398 |  |  |  | 1 |  |
| Venerida | Veneridae | *Venerupis philippinarum* | 98.66 | JF693398 |  |  | 2 |  |  |
| Myida | Corbulidae | *Corbula amurensis* | 99.39 | KJ522938 |  |  |  |  | 2 |
| Phyllodocida | Glyceridae | *Glycera chirori* | 99.05 | HZPLY772-13 | 1 |  | 1 |  |  |
| Aphragmophora | Sagittidae | *Sagitta crassa* | 99.14 | HQ700947 |  |  | 1 |  |  |
| Calanoida | Acartiidae | *Acartia bifilosa* | 98.86 | EU599508 |  |  | 2 | 3 |  |
| Rhabditida | Rhaphidascarididae | *Hysterothylacium aduncum* | 98.13 | FJ907319 |  |  |  | 1 |  |
| Total |  |  |  |  | 11 | 58 | 56 | 47 | 59 |
| Number of species |  |  |  |  | 4 | 6 | 10 | 8 | 12 |

Through nesting the ‘‘unknown’’ clones within a clade comprising of members of a single taxon in the neighbor-joining tree, three crustacean orders were identified: Mysida with 13 samples that belong to Mysidae; Amphipoda with one sample; and Decapoda with all remaining matches. Among the Decapoda, one specimen matched the genus Austinogebia, one matched to Hippolytidae, one matched to Porcellanidae and three matched to Varunidae (Fig 4).

**Fig 4.**
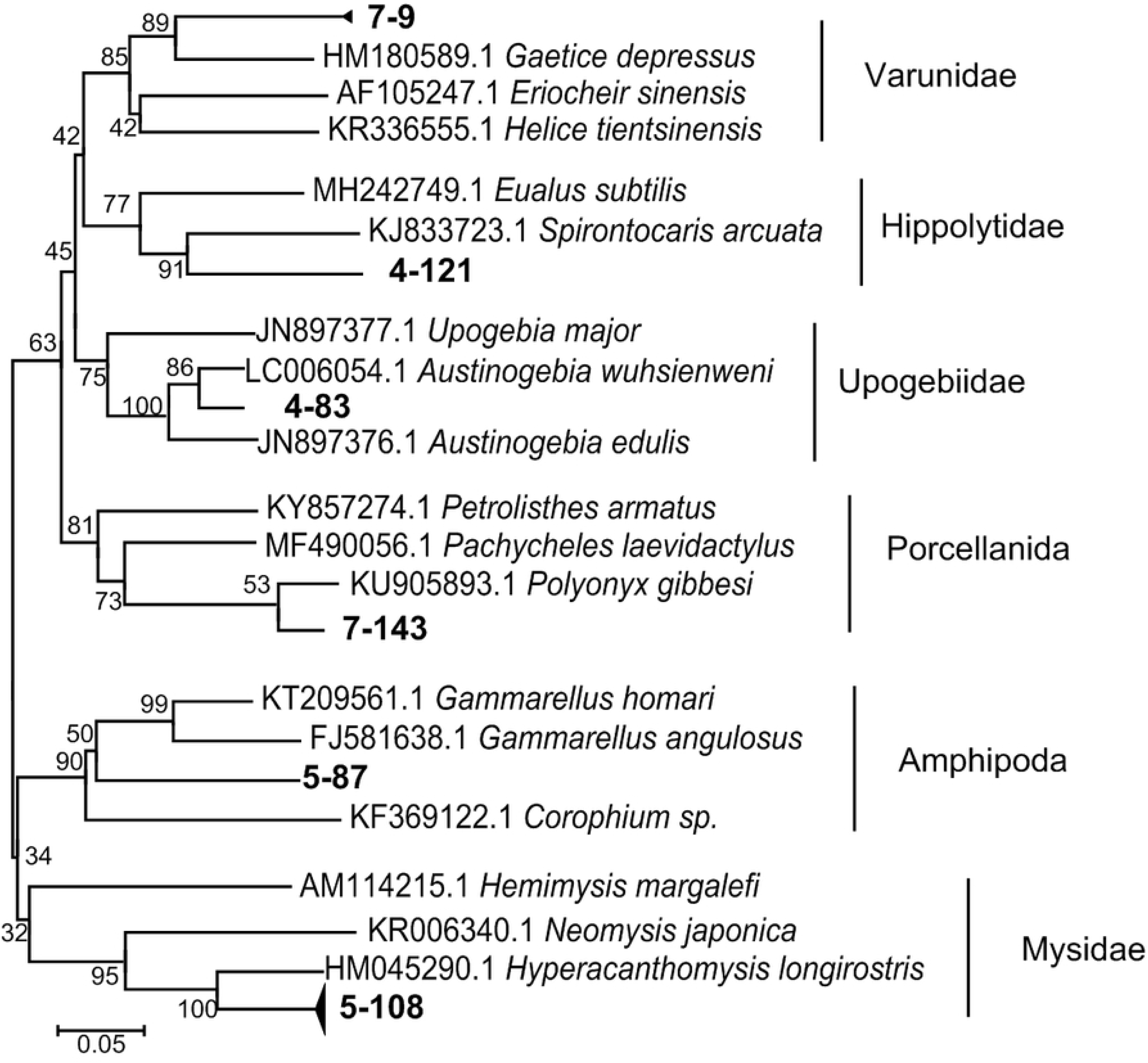
Neighbor-joining tree for 6 clades representing crustaceans in the stomach contents of the mantis shrimp. Each clade comprises of members of a single taxon and the ‘‘unknown’’ clones (bold font) were nested within a clade.

In summarising, prey detected in *O. oratoria* consisted mainly of crustaceans that accounted for 71.86 % of the clones detected, 16.02% corresponded to fishes, 8.23% corresponded to mollusks and the remaining 3.90% corresponded to other marine organisms. Fifteen orders were identified in *O. oratoria* stomachs and the number of diet species tended to increase from April to July. Six orders, each of which constituted more than 2% of the sequencing reads, accounted for 92.21% of the clones (Fig 5). Stomatopoda accounted for the highest proportion, 61.90%, followed by Gobiiformes (14.72%) and Mysida (5.63%).

**Fig 5.**
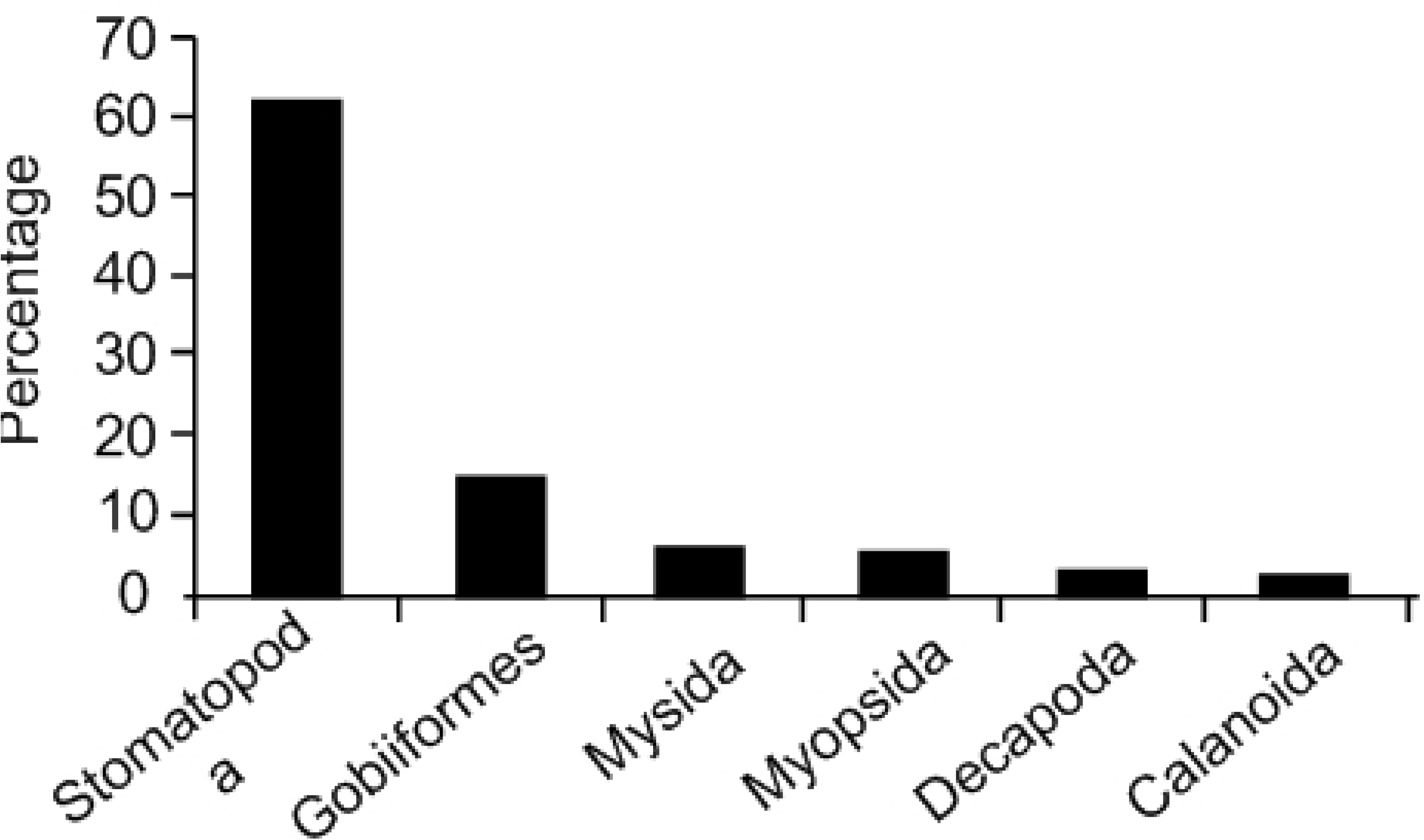
Rank abundance of orders of prey organisms detected in the stomach of O. oratoria. The y-axis shows the percentage of total reads that each taxonomic order contributed. The x-axis shows the prey orders that constituted more than 2% of the sequencing reads.

The cumulative prey curve shows that 207 stomachs were adequate to describe the diet of this species. The slopes of the saturation curves rapidly approached the asymptotes, indicating that sufficient sequencing reads were generated to capture major prey items and trends towards capturing full taxon richness (Fig 6).

**Fig 6.**
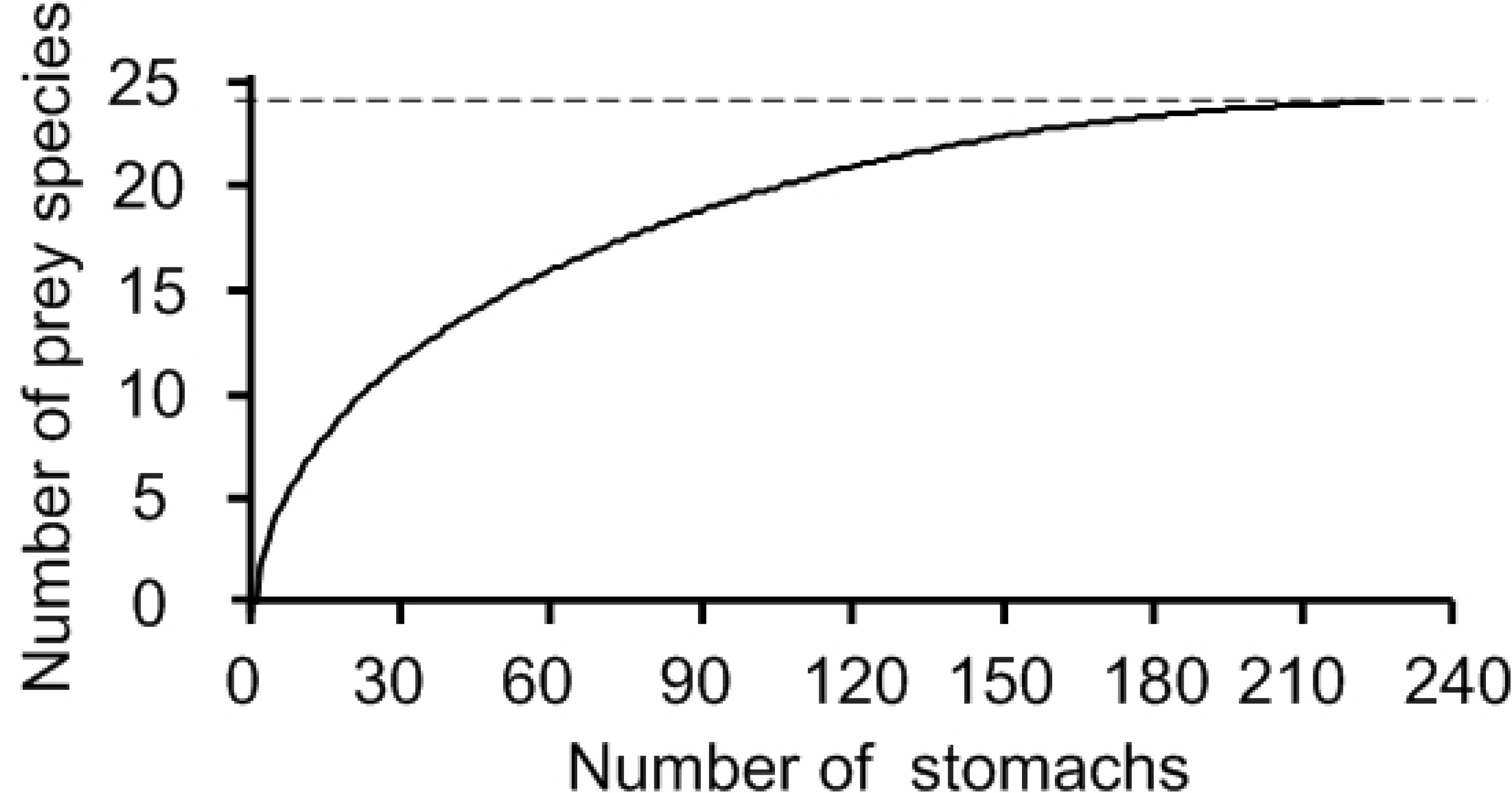
The species-accumulation curves of diet detected in 207 mantis shrimp stomachs. The asymptote represents 24 taxa.

### Cannibalism

When considering the importance of these groups in the diet of *O. oratoria*, it is remarkable that its own species was the most common prey species, detected in 143 out of 207 stomachs, accounting for 69.08% (Table 2). This study confirmed that the mantis shrimp *O. oratoria* was a cannibalistic predator in Bohai Bay. Although significance test of regression coefficients of linear (R2=0.469, P=0.201) and curvilinear regression (R^2^=0.560, P=0.167) were not significant, the degree of cannibalism decreased with an increase in diet species (Fig 7).

**Fig 7.**
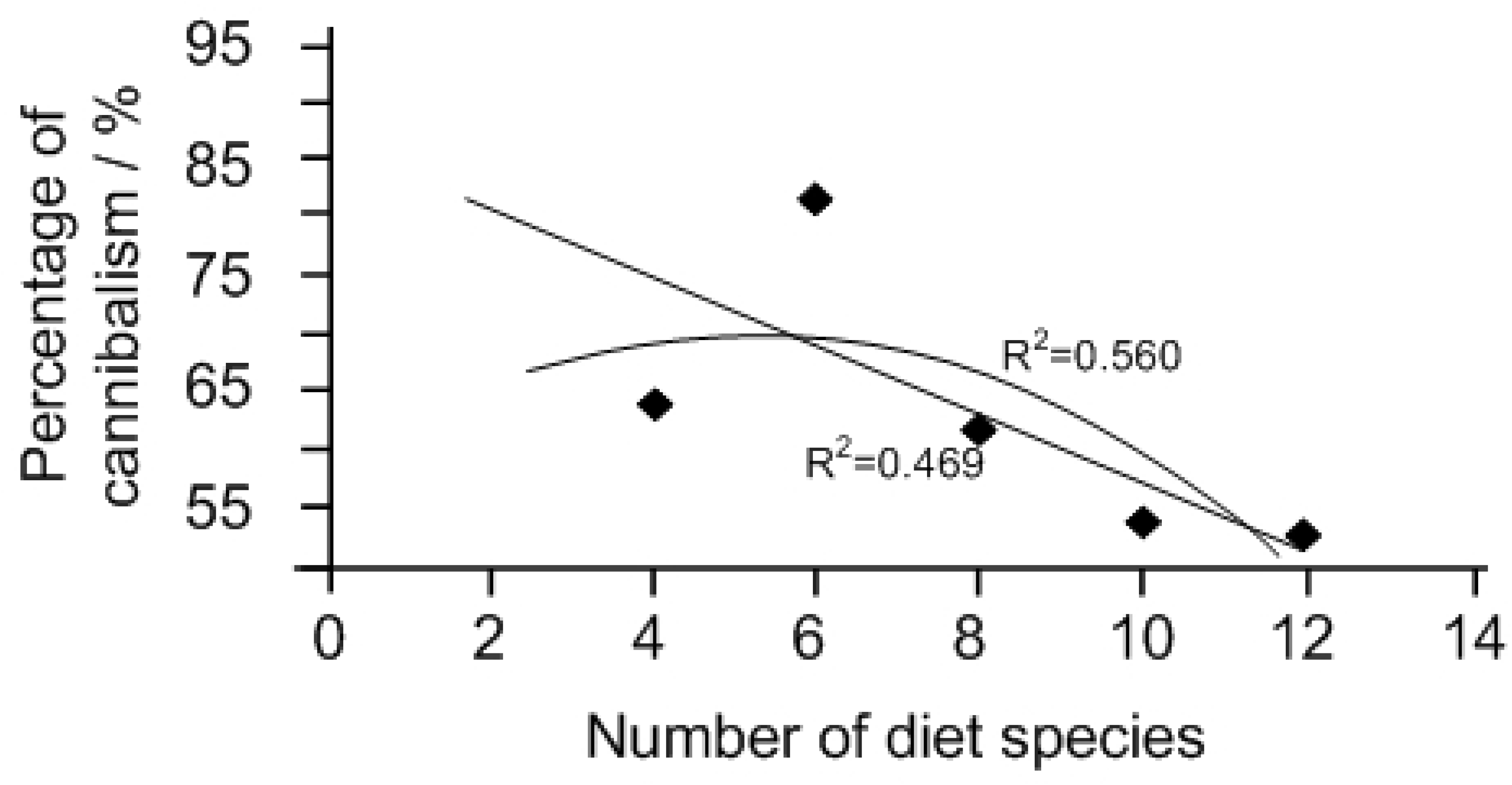
Relationship between cannibalism events and diet species; data were counted once a month.

## Discussion

Although the first peak of the gonadosomatic index in *O. oratoria* and spawning of large females were observed in spring [4], the first maturity fastigium of *O. oratoria* was found in mid-to-late May in northern China because of the overexploitation of larger individuals [1, 4–7], where a peak in lipid and protein levels in ovaries was observed in May [1]. The overexploitation of larger individuals also delayed the appearance of *O. oratoria* pseudozoea [6]. In this study, we were interested in the five months from March to July, when mantis shrimps attain gonadal maturity and then spawn. It is well known that the nutritional status of crustacean broodstock can markedly affect ovarian maturation and reproductive performance, as well as egg and offspring quality [28, 29]. Thus, investigation of the change mechanisms of the feeding strategy due to the conversion of nutrients in the body during this special period is of more research value.

An increase in the biosynthesis of various proteins, including hormones, enzymes, and lipoproteins, is involved in gonadal maturation [1, 30]. Mantis shrimps produce large numbers of yolk-laden eggs, where vitellogenesis, the process of yolk formation, is central to oogenesis [1]. It was generally thought that because of the fast energy supplement for gametogenesis from recently ingested energy, especially for females, there was an obvious peak in the feeding intensity during gonadal maturity at the end of spring [7, 8]. However, the results showed that no significant differences were found in the visual fullness index and in the FWI, suggesting that the feeding activity of *O. oratoria* was consistent between sexes and across the months and there was no obvious peak in feeding intensity. Yan et al (2017) [1] found that the beginning of reproduction of *O. oratoria* was related to reproductive effort, defined as the proportion of body energy transferred to reproduction. Their research results showed that, in both sexes, lipid contents and protein levels in the hepatopancreas and muscle decreased before May in accordance with the peak in lipid contents and protein levels in the gonads, suggesting *O. oratoria* was a conservative species, whose energy for gametogenesis comes from substrates stored in various organs and tissues (muscle, digestive gland, and mantle) through feeding prior to gametogenesis [31]. Mobilization of energy from the hepatopancreas to the gonads during periods of high energy demand is also found in other species of crustaceans [32, 33]. Compared to the opportunistic species whose energy for gametogenesis comes from recently ingested energy, the feeding intensity of conservative species tends to be stable. In view of these research conclusions, it is reasonable that the feeding activity of *O. oratoria* was consistent between sexes and no obvious peak in feeding intensity occurred prior to gonadal maturity.

Mantis shrimp are carnivorous and active predators and cannibalism occurs when large adults feed on small individuals [9]. The pleopods remains of mantis shrimps were frequently identified from stomachs and cannibalism occurred frequently (69.08%) in the result from this study and was much higher than that of the previous studies. The diets of *O. oratoria* in open seas, the Huanghai Sea and the Donghai Sea, were studied and it was found that cannibalism occurred incidentally at average value of 2.55% and 1.1% respectively [7, 8]. Hamano and Matsuura (1986) [9] found cannibalism occurred in only 0.7% of individuals when they studied the food habits of the mantis shrimp in Hakata Bay, Japan. The frequency of occurrence of mantis shrimp in stomachs remains is so high and the disparities between this study area and other areas are so great that it is necessary to think of cannibalism as a significant feeding strategy in the Tianjin coastal zone of Bohai Bay. It is thus assumed that cannibalism is part of a population energy storage strategy that enables mantis shrimps populations to react environmental conditions by reducing their numbers [34, 35].

Cannibalism is mainly and frequently a response to density or food in the field [34, 35]. The mechanism of density affecting cannibalism is that when population density increases, the territories must decrease and subsequently the frequency of intra-specific encounters and the rate of cannibalism increases [35, 36]. However, dramatic increases in the demand for stomatopods has recently led to overfishing and in the Bohai Bay, the mantis shrimp has been one of main fishing targets for crustacean fishing and is heavily caught by bottom-trawl and trammel nets. The wild stocks of *O. oratoria* have been seriously damaged [1, 5–7]. Its annual catches have decreased in the last ten years, from 1769 tons in 2007 to 510 tons in 2017 with a 71.17% drop (Fig 8) [3]. This means that the per capita area increased by nearly 2.5 times in the past 10 years and it is thus thought the density has limited influence on cannibalism in the study area.

**Fig 8.**
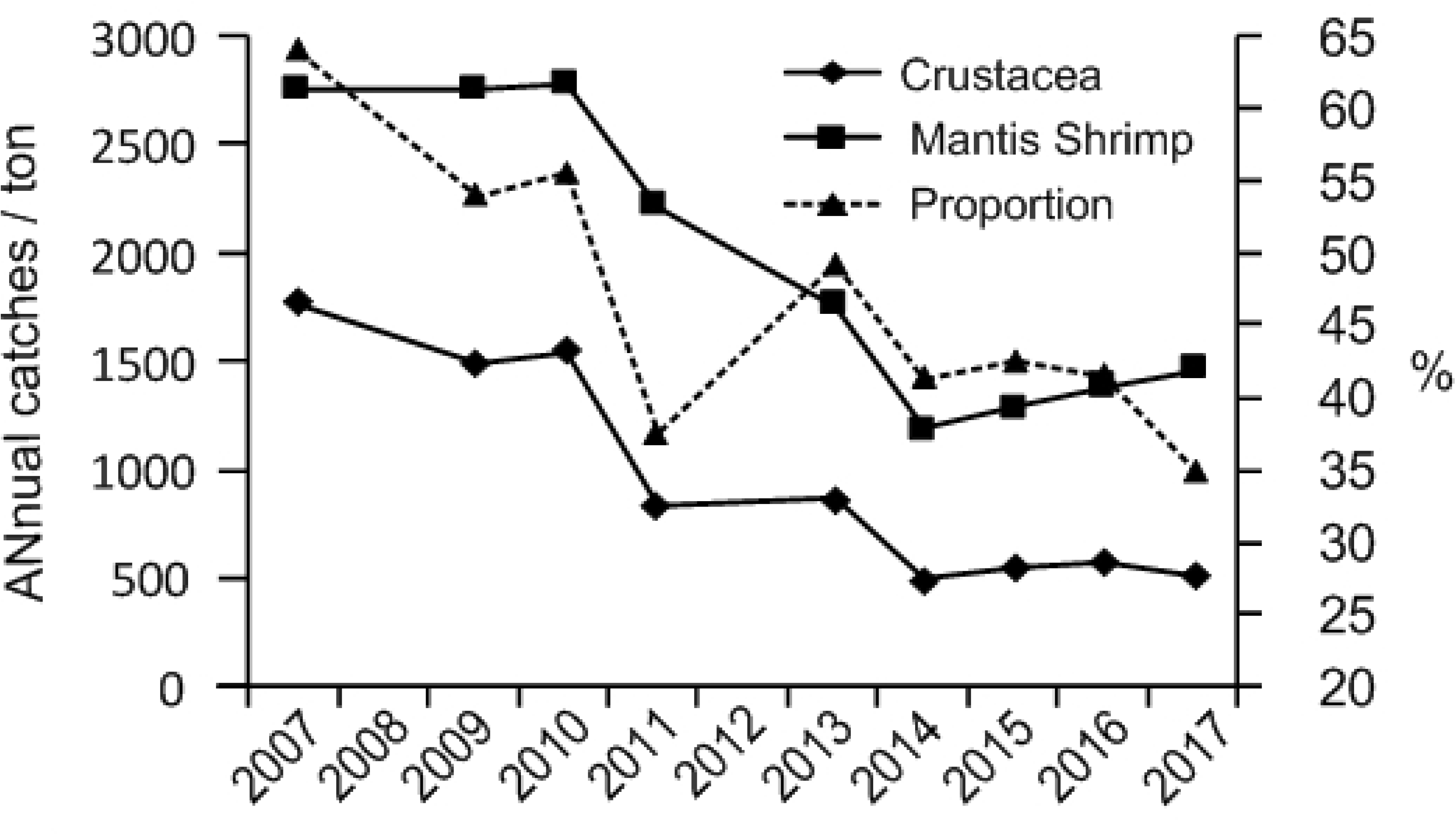
The annual catches of mantis shrimp *O. oratoria* and their proportion in crustacean catches from 2007 to 2017.

Starvation obviously increases cannibalistic tendencies [35, 37]. The mean FWI for each month ranged from 3.04 to 4.68 in this study, which is smaller than that for other waters, where the range was from 4.64 to 9.95 [7]. Additionally, the percentage of empty stomachs ranged from 28% to 65% from April to July, and was significantly greater than those of Sheng et al (2009) [7], from 4% to 20% and Xu et al (1996) [8], from 9.7% to 49%. All comparisons indicate that the mantis shrimp *O. oratoria* in the Tianjin coastal zone of Bohai Bay was suffering some degree of starvation. Under starvation, larger specimens feed on smaller conspecifics for direct food supply [34, 36] and cannibalism could provide the necessary mortality to stabilize a population during adverse conditions. A starving population with a high cannibalistic rate could have a greater chance to have an environment of sufficient production and secures reproduction [35, 38]. Moreover, cannibalism can also provide a mechanism for survival of at least parts of a population [39] as it reduces competition for the limited resources [35, 37]. The population can access lower trophic levels with the indirect extension of the food size spectrum [34] when smaller conspecifics are consumed [36].

Cannibalistic behavior has been suggested to be an indicator of limited food availability [34, 35]. The percentage of cannibalism decreased with the increase in diet species in this study (Fig 7). The study area had considerably lower biodiversity and abundance levels of macrobenthos than two other sites, the Jiaozhou Bay and the Zhoushan area of Donghai Sea [7, 8], where the diets of mantis shrimps were studied. The survey data of the macrobenthic community were obtained as closely as possible to the times when the mantis shrimps were sampled in previous studies, as listed in Table 3. The Tianjin coastal zone of Bohai Bay has experienced rapid economic and technological development and in recent years, it has also been subjected to intensive offshore exploration for production of natural gas and petroleum reserves. In addition, each year approximates 50% of China’s total maritime discharge of pollutants are brought into the sea by the river runoff. The warning of “dead sea” had drawn great concern to the Chinese government and agency [40]. The ecological environment has been affected by anthropogenic activities and these habitat changes have had a significant impact on macrobenthic communities [16, 40]. The number of macrobenthic species in this area has obviously declined compared with the historical data in 1980s, from 104.5 species to 29 species with a 72.24% species drop [40]. The average abundance of macrofauna was dramatically lower than it was ten years ago with an 86.07% drop (Table 3), and the dominant species in Bohai Bay showed a miniaturization trend where the traditionally dominant large-sized species were replaced by small-sized species, such as polychaetes and crustaceans[41, 42]. All of these changes possibly made the food for the mantis shrimp *O. oratoria* more scarce. However, there is a need for more studies to reach conclusions on how the impact of habitat changes the feeding habits of commercially important species, such as mantis shrimp.

**Table 3.** Number of species and average abundance of macrobenthos at three sites where the feeding behavior of mantis shrimp *O. oratoria* was studied. The survey data of the macrobenthic community were obtained as closely as possible to the times when mantis shrimp was sampled in previous studies.

| Sea area | Year | season | Number<br>of<br>species | Abundance<br>(ind/m<sup>2</sup>) | Sources |
| --- | --- | --- | --- | --- | --- |
| Tianjin<br>coastal zone<br>of Bohai<br>Bay | 2004 | Summer | 29 | 402.8 | [43] |
|  | 2007 | Summer | 36 | 120.27 |  |
|  | 2014 | Summer and<br>autumn | 29 | 56.07 | [40] |
| Jiaozhou<br>Bay | 2015-201<br>7 | Except<br>winter | 191 | 459.4 | [44] |
| Zhoushan<br>area of<br>Donghai Sea | 2009 | Spring | 84 | 208.5 | [45] |

The results showing that feeding activity of *O. oratoria*, a conservative species, are consistent over the months provides a reference for artificial breeding of this commercially important species that except for maintaining their normal food, the accessional diet supplied to the females bloodstocks is not a great concern because of the short culture time between collection from the wild and spawning. However what should be taken into account is that the high stocking density of broodstock wound likely cause stress responses because the frequently cannibalistic behavior in the wild makes *O. oratoria* extremely sensitive to density and the stress responses would degenerate the gonads [46]. The results also provide the trophic relationship information for fishery management and restoration of biodiversity and the abundance level of macrobenthos in Bohai Bay can improve feeding conditions for mantis shrimp, then increasing its production, because the mantis shrimp could minimize their energy expenditure when they captured and handled smaller organisms, rather than preying on their own kind with harder exoskeletons and a more formidable defensive weapon, the raptorial claw [9].

## Supporting information

**S1 File. The representative sequences detected in stomach contents.**

(PDF)

## Acknowledgments

We are indebted to deputy senior engineer Zhen-Guo Zhang and Zhang Luo for their kind assistance in molecular experiments and to intermediate engineer J Yu for her kind assistance in specimen collection. We also thank American Journal Experts (AJE) for English language editing.

